# Quantitative stable isotope probing to measure *in situ* growth of protist populations

**DOI:** 10.64898/2026.06.04.730198

**Authors:** Rebecca L. Mau, Michaela Hayer, Paul Dijkstra, Stefan Geisen, Bruce A. Hungate, Egbert Schwartz

## Abstract

Soil protists shape microbial food webs and nutrient cycling, yet methods for measuring their population growth in soil have lagged behind the taxonomic resolution available from 18S rRNA gene sequencing. Microscopy-based approaches can estimate abundance and growth, but with limited taxonomic resolution. Here, we tested whether quantitative stable isotope probing (qSIP) can provide reproducible, sequencing-resolved growth measurements for soil protists. We incubated soil with natural-abundance or ¹ O-labeled water and measured taxon-specific ^18^O enrichment in DNA using three common 18S rRNA gene primer sets. ^18^O enrichment values were positively correlated across datasets, with relationships closest to 1:1 after poorly resolved taxonomic assignments were excluded, indicating that qSIP provides reproducible population-level growth signals across marker choices. We then compared 18S amplicon profiles from unfractionated DNA with qSIP-derived growth measurements across a soil moisture gradient. Amplicon profiles showed small shifts in relative abundances of major protist groups, whereas qSIP revealed a large moisture response in the growing community: 7 ASVs were growing at 20% field capacity compared with 143 at 80% field capacity, representing 1.6% and 63.3% of total protist relative abundance, respectively. Average growth was <1% day□^1^ in the two driest treatments, increasing to 2.1% day□^1^ at 60% and 5.6% day□^1^ at 80% field capacity; among growing taxa, rates averaged 8.7% and 8.3% day□^1^ in the two wetter treatments. By pairing taxonomic resolution with isotope-based growth estimates, qSIP with ^18^O-H_2_O moves soil protist ecology toward quantitative population dynamics: identifying which taxa grow, how fast, and how growth responds to the environment.

## Introduction

Soil protists are taxonomically diverse eukaryotes that influence terrestrial carbon and nutrient cycling as primary producers, consumers, predators, and parasites. In soil food webs, protist consumers can alter bacterial and fungal abundance, community composition, decomposition, plant health, and nitrogen release through predation [1–6]. Quantifying these effects in soil remains difficult because soil is spatially heterogeneous and opaque, and because many protists shift rapidly among growing, active, and dormant states.

Microscopy-based approaches, including most probable number assays, liquid aliquot methods, and direct counts, have provided important measurements of protist abundance, biomass, morphology, and behavior [15–20]. Their strengths are direct observation and absolute counts. Their limitations include coarse taxonomic resolution, culture bias, and potential stimulation of encysted organisms during liquid extraction or incubation [21–23].

High-throughput 18S rRNA gene amplicon sequencing has become a central tool for soil protist ecology because it provides broad taxonomic coverage and reveals diversity that microscopy cannot resolve [27–29]. Amplicon surveys also inherit biases from DNA extraction, primer choice, PCR conditions, reference databases, and relic DNA [30–35]. While DNA inventories describe which taxa are represented in a sample, population dynamics require measurements of which taxa are making new cells. RNA-based approaches provide valuable metabolic information over short time scales, although rRNA abundance can be a poor proxy for activity and mRNA is difficult to link to taxonomic identity in complex soil communities [36, 37]. Stable isotope probing (SIP) was developed to link microbial identity with metabolic processes *in situ* [38], and quantitative SIP (qSIP) extends this approach by estimating isotope enrichment and population growth for individual taxa [39]. With H_2_^18^O-qSIP, taxa that synthesize DNA during incubation incorporate ^18^O from water into new DNA, shifting their buoyant density in a way that can be quantified after isopycnic centrifugation and amplicon sequencing [40–43].

The value of qSIP is that isotope enrichment is a quantitative population-level measurement. For each taxon that passes filtering criteria, qSIP returns an estimate of isotope incorporation that can be translated into the fraction of new DNA and, with an explicit growth model, population growth rate. These values enable comparing the magnitude of responses among taxa, treatments, and time points. This places qSIP in a broader quantitative isotope tradition in which isotope values estimate process rates, like ¹□N isotope pool-dilution measurements of gross nitrogen mineralization, microbial immobilization, and nitrification in intact soils [49, 50].

Moisture provides a biologically interpretable test system for qSIP measurements of protist growth. Most soil protists require connected water-filled pores for mobility, prey encounter, and feeding [51, 52], and many studies have reported positive relationships between soil water and protist abundance following rain events, across seasons, and in laboratory wet-up experiments [1, 15, 20, 53–55]. Molecular growth measurements can extend this literature by identifying which protist populations synthesize new DNA under specific moisture conditions and by estimating the magnitude of their population growth during soil incubation.

We used H_2_^18^O-qSIP to measure taxon-specific DNA replication and population growth in soil protist communities. Our first objective was to test whether qSIP produced reproducible ^18^O enrichment values across three commonly used 18S rRNA gene primer sets: two universal primer pairs that target different regions of the 18S rRNA gene (V9, [57]; V4, [58]) and one primer set that targets Cercozoa ([59]). Our second objective was to compare unfractionated 18S amplicon profiles with qSIP-derived growth measurements across a soil moisture gradient. We expected increasing soil moisture to increase the richness, relative abundance, and growth rates of growing protist populations because connected water films should increase protist mobility and prey access. We also expected amplicon profiles and qSIP measurements to produce different views of the community because DNA inventories reflect the total recovered DNA pool, whereas qSIP identifies the taxa making new DNA during incubation.

## Materials and Methods

### Experimental setup

Soil (0-10 cm) was collected from the Hopland Research and Extension Center in Hopland, CA in August 2021. Five g field-dry soil (1.8% gravimetric water content) was weighed into 15 mL Falcon tubes and water was added to generate soil moisture treatments at four different field capacities: 20%, 40%, 60%, 80%, equivalent to 7.2%, 14.4%, 21.6% and 28.8% gravimetric water content, respectively. Half of the samples (*n*=5) received natural abundance deionized water, and the other half received 97 atom% ^18^O-labeled water. Soils were incubated in the dark at room temperature for five days and then frozen at −20°C. DNA was extracted from 0.5 g soil in triplicate using a Qiagen DNeasy PowerSoil kit following the manufacturer’s protocol and was then pooled together. Five µg DNA was spun in a cesium chloride solution consisting of 3.65 mL saturated cesium chloride and approximately 950 μL gradient buffer solution (200mM Tris pH 8, 200mM potassium chloride, 2mM EDTA). Samples were spun at 104,114 x *g* for 72 hours in a Beckman benchtop ultracentrifuge (Beckman Coulter, Brea, CA) using a TLA-110 rotor with 4.7 mL ultracentrifuge tubes. Samples were fractionated into ∼21 - 200 µL aliquots and DNA was purified using an isopropanol precipitation method [41]. Density standards were added to the centrifugation tubes prior to ultracentrifugation and were detected in sample-fractions using qPCR, allowing for the calculation of the density of each sample-fraction [43, 60]. Because the buoyant density of a taxon’s DNA, determined by the GC content of DNA [61], is unlikely to change over a five-day incubation and because of the costly and laborious process, only natural abundance samples from the 80% field capacity treatment were processed (*n*=5) to represent the buoyant density of a taxon’s natural abundance DNA in all soil moisture treatments.

To compare ^18^O enrichment values calculated from datasets targeting different 18S rRNA gene regions, qPCR was performed on all sample-fractions from the 80% field capacity treatment using primers that targeted the V9 (1380F/1510R primer pair;, V4 (TAReukFWD1/REV3 primer pair;, and Cercozoa-specific V4 18S rRNA gene regions (S616F_Cerco;S616F_Eocer/S963R_Cerco primers;. Master mixes and thermocycling conditions for each primer pair are presented in Table 1. qPCR was performed on all sample-fractions from the 20%, 40%, 60% field capacity treatments using primers targeting the V4 18S rRNA gene region only (TAReukFWD1/REV3 primer pair).

**Table 1.**
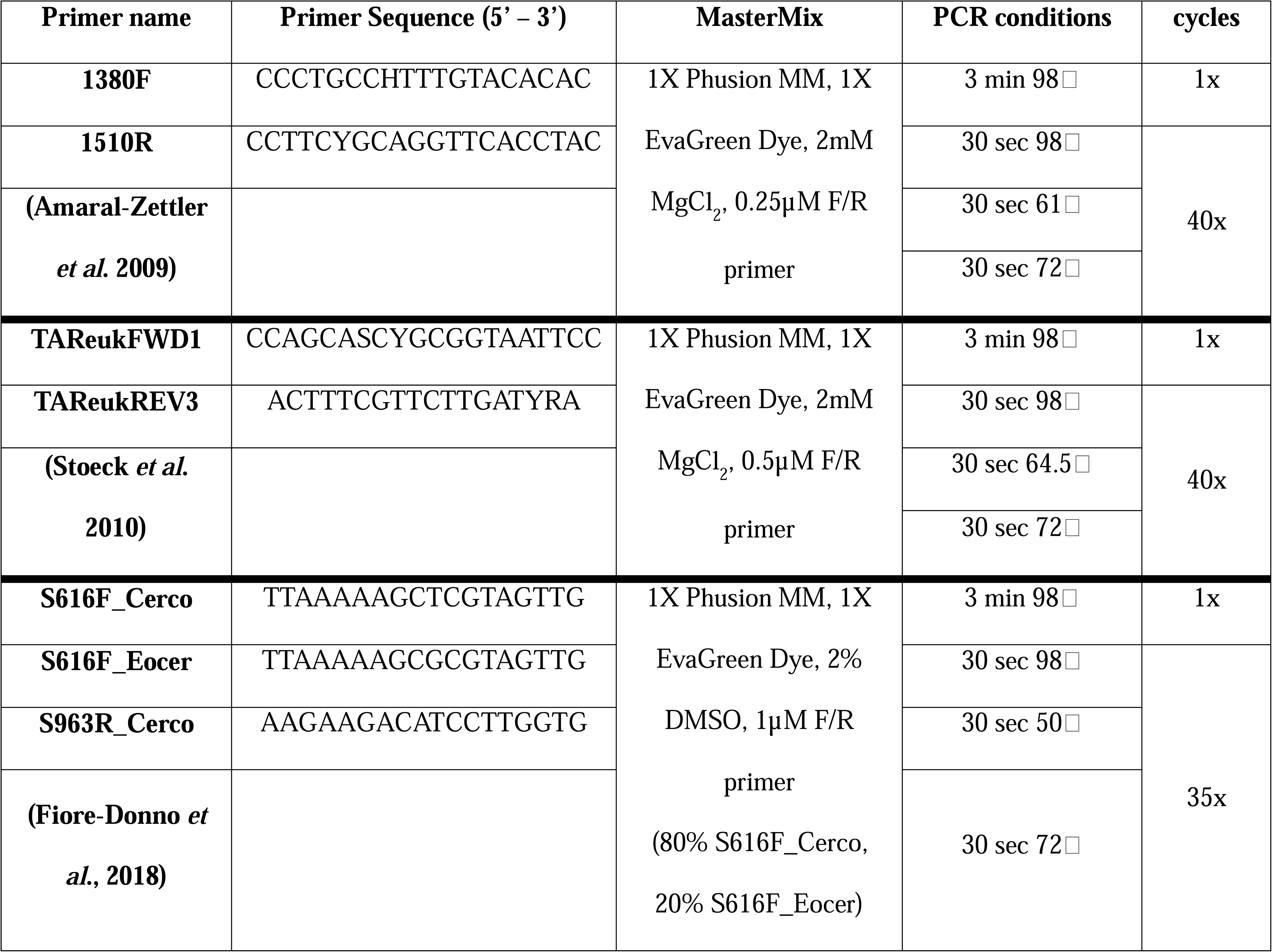
The PCR conditions and master mix compositions used in this study for the three primer pairs used to target the V9 18S rRNA gene region (1380F/1510R), V4 18S rRNA gene region (TAReukFWD3/REV1), and the Cercozoa-specific V4 18S rRNA gene region (S616F_Cerco,S616F_Eocer/S963R).

Sequencing libraries were then prepared using the same samples and primer pairs used for qPCR that targeted different 18S rRNA loci using sample-fractions in the density range of 1.66 – 1.76 g mL^−1^. A two-step PCR approach was used to prepare sequencing libraries. The first PCR was performed with the same mastermix and thermocycler conditions as for qPCR (Table 1) except all primer pairs had universal tails attached (UT-F: 5’-ACCCAACTGAATGGAGC-3’, UT-R: 5’-ACGCACTTGACTTGTCTTC-3’;), no EvaGreen dye, and was only amplified for 20 cycles. A second PCR was then conducted using amplicons from the initial PCR to attach a unique 8 bp barcode as well as the Illumina flow cell adapter tails (P5/P7) to samples in a mastermix that contained 1X Phusion Green master mix (Fisher Scientific Inc.) and 1µM F/R barcoded primer. PCR products were purified using AMPure magnetic beads (Beckman Coulter, Brea, CA) and quantified using a BioTek plate reader and the Quant-iT DNA quantification kit (Thermo Fisher Scientific, Hampton, NH). Samples from each primer pair were pooled together at equimolar concentrations and sequenced on an Illumina MiSeq instrument (Illumina, San Diego, CA). The library targeting the V9 18S rRNA gene region was sequenced using a MiSeq 2 x 150 cycle v2 kit, the library targeting the V4 18S rRNA gene region was sequenced using a MiSeq 2 x 300 cycle v3 kit, and the library targeting the Cercozoa-specific V4 18S rRNA gene region was sequenced using a MiSeq 2 x 250 cycle v2 kit. DNA that was not spun and fractionated from all samples were also sequenced using primers targeting the V4 18S rRNA gene regions, prepared as described above. Sequences have been deposited into NCBI Sequencing Read Archive (SRA) under BioProject ID PRJNA1473953.

### Bioinformatics

Sequences were processed using the QIIME2 bioinformatics platform (version 2024.2 [64]). Demultiplexed sequencing reads were imported into QIIME2 and primers were trimmed off using the cutadapt plugin [65]. Sequences were trimmed to 120 bp for the V9 dataset and 230 bp for the Cercozoa specific dataset and reads were merged during the denoising step using the DADA2 plugin (‘denoise-paired’ command; [66]). For the V4 dataset, only the forward reads trimmed to 280 bp were used (‘denoise-single’ command). Sequences from the libraries were then assigned taxonomy using the ‘feature-classifier’ command [67] and the PR^2^ reference database (5.0.0; [68]) with the percent identity parameter set to 0.9 to ensure high confidence in taxonomic assignments. Sequences that were not assigned to protist taxonomy were excluded from subsequent analyses. These assignments included: Bacteria; (and lower), Archaea;_, Eukaryota;_, Eukaryota;Archaeplastida;_, Eukaryota;Archaeplastida;Streptophyta;_(and lower), Eukaryota;Obazoa;_, Eukaryota;Obazoa;Opisthokonta;_, Eukaryota;Obazoa;Opisthokonta;Fungi;_ (and lower), Eukaryota;Obazoa;Opisthokonta;Metazoa;_(and lower), and Unassigned;_

### 18O Excess Atom Fraction (EAF) and growth rate calculations

ASV-specific ^18^O EAF values were calculated following Hungate et al. (2015) using the qSIP2 R package [69] for ASVs that were present in a minimum of four fractions and a minimum of three treatment replicates. Briefly, a change in the density of an organism’s DNA (Z*_i_*) was measured by calculating the difference in weighted average density (WAD; W*_i_*) of an organism’s DNA in samples incubated with 97-atom% ^18^O-labeled water (W_LAB_) and natural abundance water (W_LIGHT_):

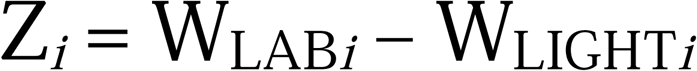

The difference in WAD is the result of the incorporation of isotopically labeled oxygen into the DNA of growing organisms [39]. WADs of taxa were calculated by first estimating the number of a taxon’s gene copies in each sample-fraction by multiplying the total number of gene copies by the relative abundance of that taxon in the sample-fraction. While we acknowledge that gene copy abundance (as estimated with qPCR) and population relative abundance (as estimated with sequencing data) were measured separately on the same sample-fractions, we assume that the use of identical master mix and PCR conditions minimized biases between the methods. The proportion of each taxon’s gene copies in a sample-fraction was then multiplied by the density of the sample-fraction and added together to get a taxon-specific WAD.

The average molecular weight of natural abundance DNA is 307.939 g mol^−1^ (M_LIGHT_; [39]). Maximum labeling of a DNA molecule occurs when all natural abundance oxygen atoms are replaced with ^18^O-labeled atoms, which equals an increase in molecular weight of 12.07747 g mol^−1^ [39]. Therefore, the theoretical maximum molecular weight of DNA (M_HEAVYMAX_), which would equal 100 atom percent ^18^O, or 1 atom fraction ^18^O, is:

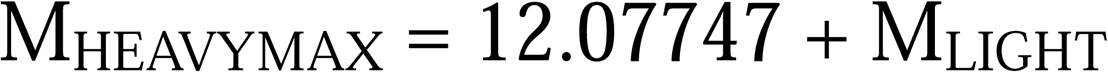

To calculate the excess atom fraction (EAF) of ^18^O per taxon, we first calculate the molecular weight of DNA per taxon (M_LAB*i*_) as the change in density of samples incubated with 97 atom percent ^18^O water and natural abundance water (Z*_i_*) relative to the density of the natural abundance sample (W_LIGHT*i*_) multiplied by the molecular weight of the natural abundance sample (M_LIGHT*i*_):

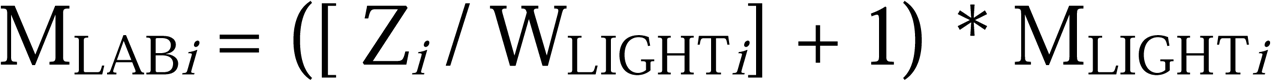

The ^18^O EAF per taxon (A_OXYGEN*i*_) is then calculated as:

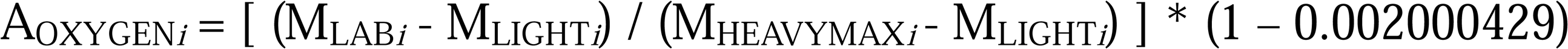

which accounts for the background abundance of ^18^O in the environment. ^18^O EAFs were then adjusted by the soil water ^18^O atom % in each treatment since the proportion of ^18^O water to natural abundance water present prior to incubation differed between soil moisture treatments.

We then assigned ASVs to a functional class using EukFunc [70] in R, which included phototrophs, parasite symbiotrophs, and predators. Out of 346 ASVs that passed the qSIP filtering criteria, 45 could not be assigned a functional class and were excluded from further analyses. Because oxygen in DNA can be derived from two sources, water and nutrients, phototrophs and parasites/symbionts will only incorporate ^18^O into their DNA from water whereas predators can incorporate ^18^O into their DNA from both water and from prey that incorporated ^18^O into their DNA. To allow for the direct comparison of enrichment, and thus growth, between organisms with different energy pathways, a correction was applied to predator taxa after [71]:

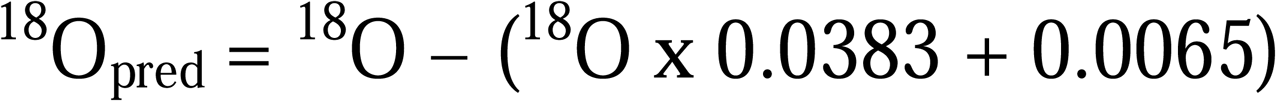

To calculate growth rates in response to soil moisture after five-days, a linear growth model was assumed and ^18^O EAF values were adjusted by the parameter *U*, which enables a more accurate estimate of growth by accounting for the maximum proportion of oxygen atoms derived from the labeled water. We used a value of 0.6 for *U* here after a sensitivity analysis of a 16S rRNA dataset [72] and data from a population that became completely enriched in this dataset (no unlabeled 18S rRNA genes at low density) were in agreement (0.6 for 16S and 0.62 for 18S).

### Statistical analyses

To compare ^18^O EAF values calculated from different primer pairs in the 80% field capacity soil moisture treatment samples, an average ^18^O EAF value was calculated from ASVs assigned to the same taxonomy. Because the different datasets used primers targeting different 18S rRNA gene regions, ASVs could not be compared directly. Average ^18^O EAF values were compared by major axis Model II linear regression analysis using R Statistical Software (v 4.3.1; R Core Team 2023) and the “lmodel2” package [73]. To test if the slope of the relationships between calculated ^18^O EAF values of shared taxa between datasets for each primer pair was significantly different from 0, a permutation test was used with 999 iterations.

Weighted average densities (WAD) of the 18S rRNA gene copies, ^18^O EAF values, and growth rates were all non-normally distributed as determined using a Shapiro-Wilk test. Therefore, the non-parametric Kruskal-Wallis test was used to determine if there were statistically significant differences in the WADs of 18S rRNA gene copies, ^18^O EAF values, and growth rates between the four soil moisture treatments. If significant values were found (p<0.05), a Dunn’s post hoc test was performed to determine which groups were significantly different from one another. Significant differences in the relative abundances of protist groups in response to soil moisture were determined with ANOVA and Tukey’s post-hoc analyses when significant. Finally, we used bootstrapping (1001 iterations with replacement) to determine the average and 90% confidence intervals of the ^18^O EAF values and growth rates.

## Results

### Taxon-specific ^18^O enrichment values were reproducible across primer sets

The V4 and V9 datasets had 35 taxonomic assignments in common, including taxa from the Alveolata (3), Amoebozoa (3), Cercozoa (15), and Stramenopiles (8) groups. There was a significant positive relationship between the ^18^O EAFs calculated from the V9 and V4 datasets from all 35 shared taxa (slope=0.603, *p*=0.008); however, the slope of this relationship became closer to 1 when taxa assigned to order or higher were excluded, which resulted in 26 taxonomic assignments in common (slope=0.751, *p*=0.001; Fig 1A). The V9 and Cercozoa-specific dataset only shared 17 taxa and the relationship between the ^18^O EAFs of these two datasets was significantly positive (slope=1.66, *p*=0.012). The slope of this relationship also became closer to 1 when data from three taxa assigned to order or higher were excluded (slope = 1.07, *p*=0.001; Fig 1B). The V4 and Cercozoa datasets shared 40 taxa and also showed a strong positive relationship of ^18^O EAF values between the two datasets (slope=0.991; *p*=0.001). Filtering out taxa assigned to order or higher resulted in 35 shared taxa with a similar slope (slope=1.01; *p*=0.001; Fig 1C).

**Figure 1.**
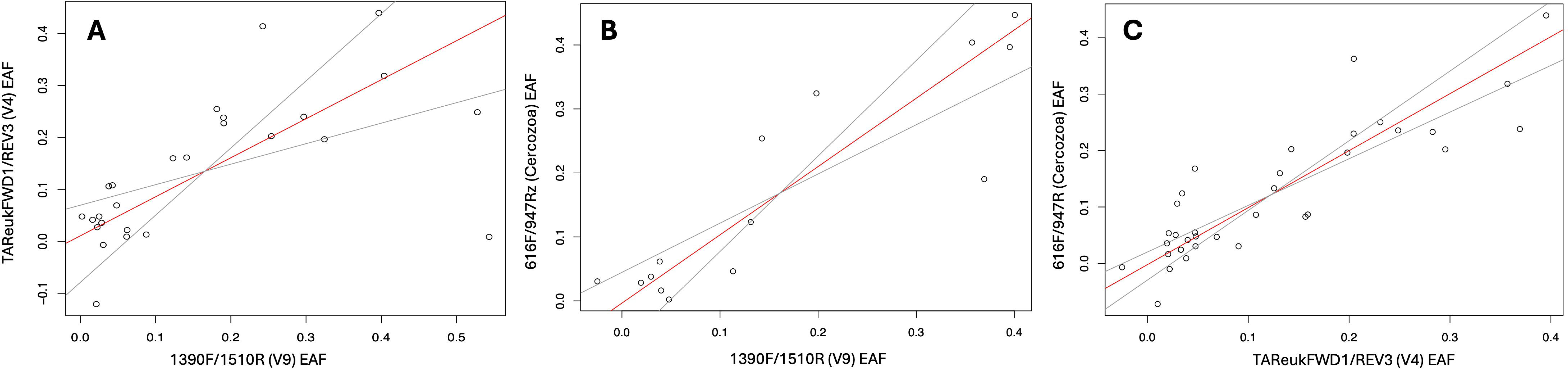
Relationships between ^18^O excess atom fractions (EAF) of taxa calculated from datasets generated using different primer pairs that targeted different 18S rRNA gene regions: V9 vs V4 18S rRNA gene regions (A), V9 vs Cercozoa-specific V4 18S rRNA gene regions (B), V4 vs Cercozoa-specific V4 18S rRNA gene regions (C). The red lines show the fitted line and the grey lines are 95% confidence interval limits.

### Soil moisture increased protist DNA replication and population growth

^18^O incorporation into the DNA of the protist community increased as soil moisture increased as seen by an increase in the proportion of protist 18S rRNA genes in the heavy fractions (∼1.715 g mL^−1^ ∼ 1.75 g mL^−1^) of samples incubated with ^18^O-water compared to samples incubated with natural abundance water (Fig 2A). Furthermore, the WAD of protist 18S rRNA genes incubated with ^18^O-water at 80% field capacity was significantly heavier (*H* = 16.0, *p* = 0.003) than the WAD of 18S rRNA genes incubated with natural abundance water and soils at the two lowest moisture treatments (1.716 ± 0.0007 g mL^−1^ vs 1.704 ± 0.0004 g mL^−1^; Fig 2B).

**Figure 2.**
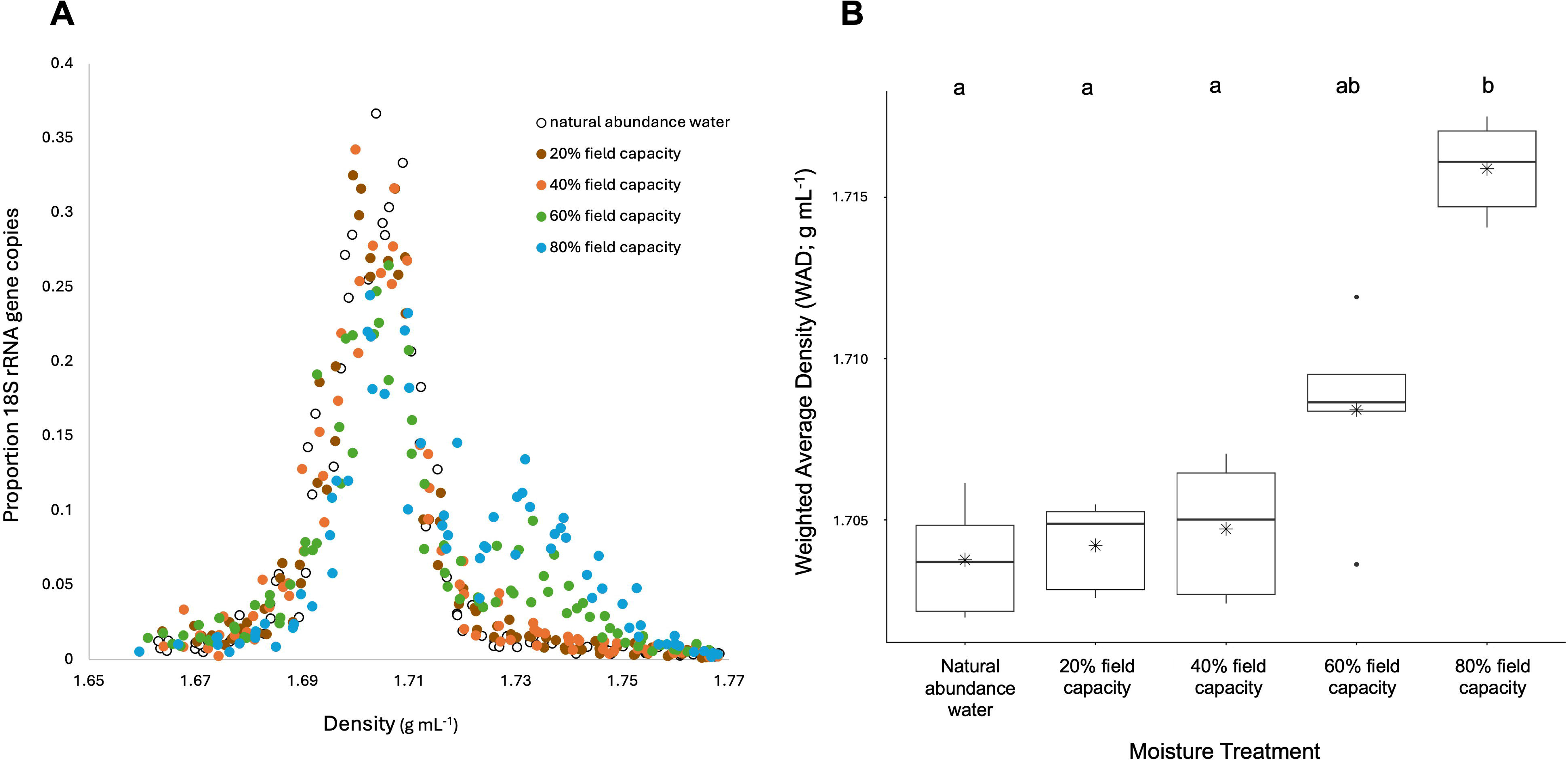
Proportion of 18S rRNA gene copies at densities between 1.66 – 1.77 g mL^−1^ (A) for samples incubated with natural abundance ^18^O-water (open circles) or 97 atom% ^18^O-water at four different moisture treatments: 20% field capacity (brown circles), 40% field capacity (orange circles), 60% field capacity (green circles), and 80% field capacity (blue circles). Panel B shows significant differences (*H* = 16.0, *p* = 0.003) in the weighted average densities (WADs) of 18S rRNA gene copies of soils at the same five treatments (natural abundance ^18^O-water, 20%, 40%, 60%, and 80% 97 atom % ^18^O-water) as determined by a Kruskal-Wallis test. Letters denote significant differences from a Dunn’s post-hoc test.

There were 1915 protist ASVs in the sequencing dataset from the unfractionated samples (i.e. traditional amplicon sequencing approach) from the four different soil moistures, representing 286 unique taxonomic assignments. Of the 1915 ASVs, 158, 191, 249, and 248 ASVs passed our qSIP filtering criteria of being present in a minimum of four fractions and three replicates in the 20%, 40%, 60%, and 80% field capacity moisture treatments, respectively. A further 29, 30, 36, and 33 ASVs were filtered out from the 20%, 40%, 60%, and 80% field capacity moisture treatments, respectively, because they were not assigned a functional class. This resulted in 65.3%, 69.0%, 77.1%, and 78.7% of the relative abundances of the total protist communities that were included in the qSIP growth rate calculations (Fig 3B) and represented 116 unique taxonomic assignments.

**Figure 3.**
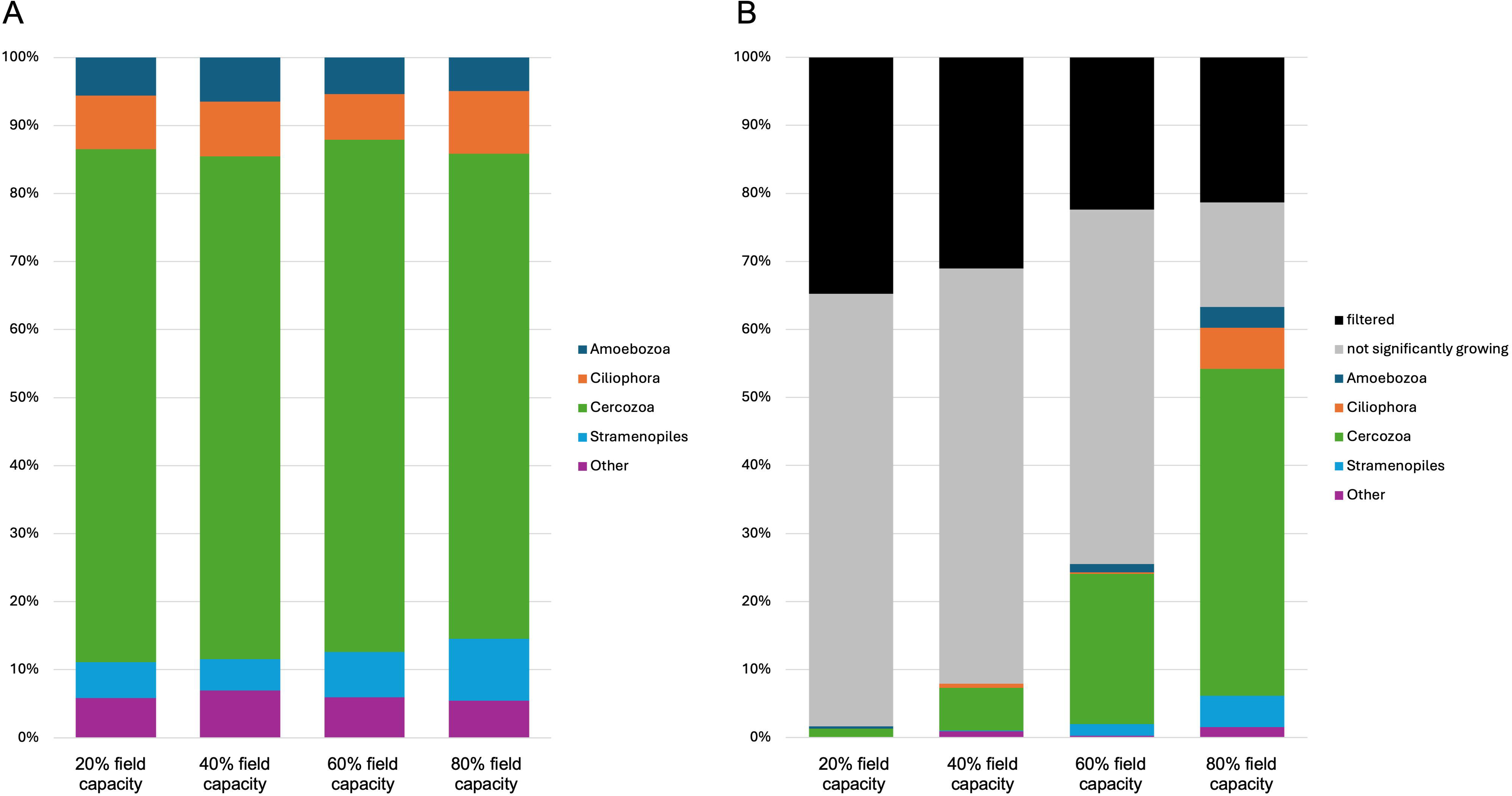
Relative abundances of protist groups for samples incubated for five days at 20%, 40%, 60%, and 80% field capacity using a traditional amplicon sequencing approach (A) or a quantitative stable isotope probing approach (B). Protist groups that made up <1% relative abundance were grouped into the “Other” category. In panel B, ASVs that were not in 3 treatment replicates and/or 4 sample fractions were filtered out and denoted as “filtered” in the figure. Taxa whose population growth 90% confidence intervals overlapped zero were considered “not significantly growing” in panel B.

The relative abundance of Amoebozoa was significantly lower in the 80% field capacity treatment compared to the 40% field capacity treatment (*F* = 3.38, *p* = 0.044) and the relative abundance of Stramenopiles was significantly higher in the 80% field capacity treatment than in the 20% or 40% field capacity treatments (*F* = 7.70, *p* = 0.002; Fig 3A). Both sequencing approaches identified Cercozoa (TSAR;Rhizaria) as the dominant group, making up an average of 74.2% ± 0.9% of the relative abundance of protists in the traditional amplicon sequencing approach (Fig 3A) and 81.1% ± 2.2% of the significantly growing protist communities (Fig 3B). Ciliates (TSAR;Alveolata;Ciliophora) were the second most abundant group at all soil moistures in the traditional amplicon data (8.0% ± 0.5%; Fig 3A) but only made up a large proportion of the growing populations in the 80% field capacity treatment (Fig 3B).

The proportion and richness of the growing protist population dramatically increased as soil moisture increased (Fig 3B). Populations of only seven ASVs were identified as significantly growing (90% confidence intervals did not cross zero) when soil moisture was at 20% field capacity (Fig 4), making up a mere 1.6% of the relative abundance of the protist community (Fig 3B). At 40%, 60%, and 80% field capacity, 15 taxa (18 ASVs), 31 taxa (63 ASVs) and 79 taxa (143 ASVs) were significantly growing, respectively (Fig 4). These taxa made up 4.9%, 25.5%, and 63.3% of the protist community relative abundance, respectively (Fig 3B).

**Figure 4.**
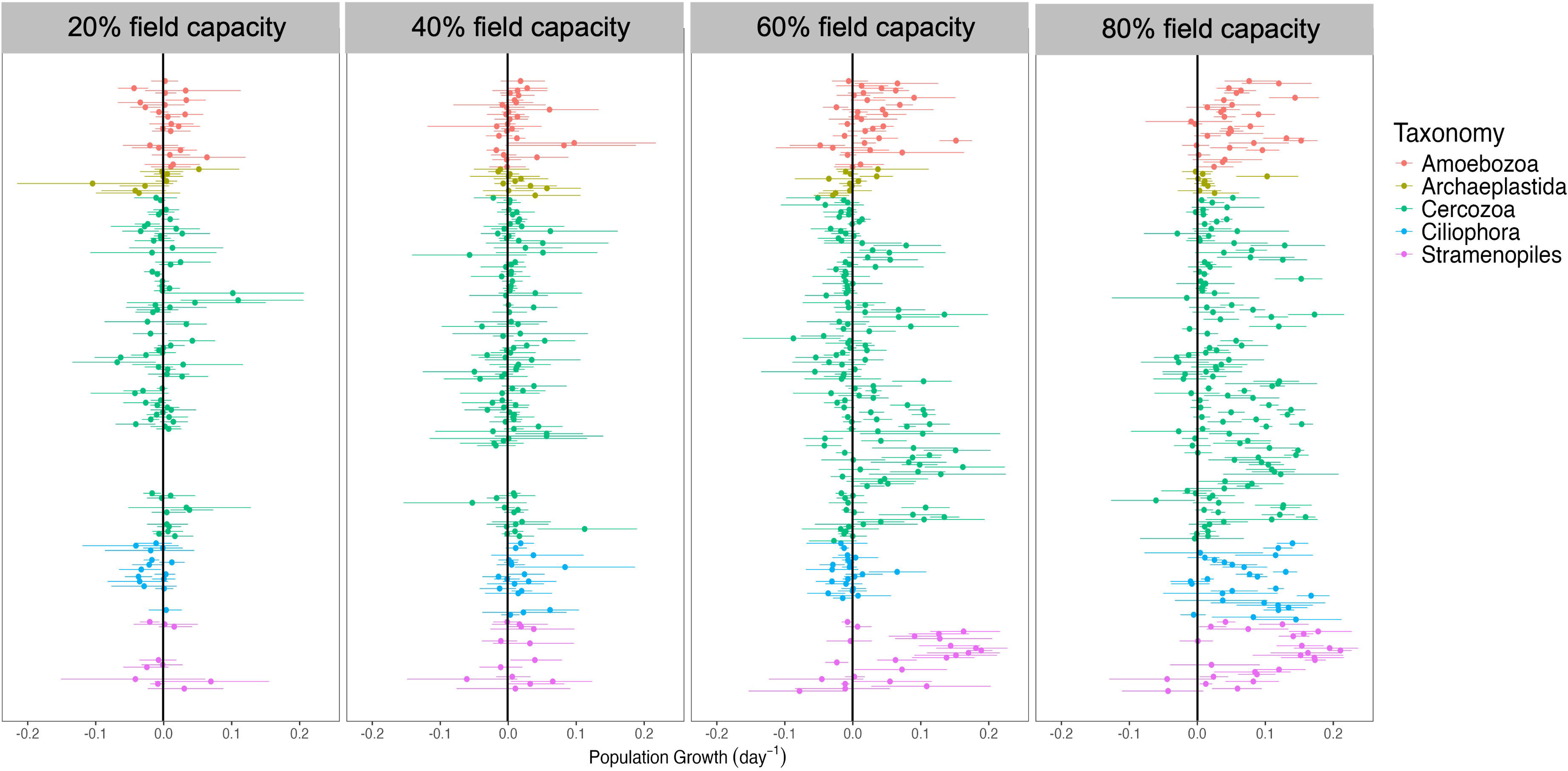
Taxon specific population growth rates (day^−1^) of protists colored by taxonomic groups in soil incubated for five days at 20%, 40%, 60%, or 80% field capacity. Points are average growth rates and lines represent 90% confidence intervals. Populations whose 90% confidence intervals overlapped zero were considered “not significantly growing.”

As hypothesized, population growth rates increased with increasing soil moisture. The average population growth rate of the total protist community was 5.6% day^−1^ at 80% field capacity, 2.1% day^−1^ at 60% field capacity, and <1% day^−1^ at 20% and 40% field capacities (*H* = 129.3, *p* < 0.0001; Fig 5A). The top two fastest growing ASV populations were found in the highest soil moisture treatments and were assigned to the Amphililaceae family (TSAR;Stramenopiles;Bigyra;Sagenista;Labyrinthulomycetes). These two ASVs were growing at an average rate of 20.3% ± 0.8% day^−1^ and 18.5% ± 0.4% day^−1^ in the 80% and 60% field capacity treatments, respectively (Supplemental Table 1). Some of the slowest growing taxa after five days were two ASVs from the *Thaumatomonas* genus (TSAR;Rhizaria;Cercozoa:Filosa-Imbricatea;Thaumatomonadida;Thaumatomonadidae; avg 0.79% day^−1^) and three Cryomonadida ASVs in the Rhogostoma-lineage (TSAR;Rhizaria;Cercozoa;Filosa-Thecofilosea; avg 1.3% day^−1^), both at 80% field capacity. While population growth rates of the total protist population were close to zero in the 20% and 40% field capacity soil moisture treatments, the average population growth rates of the growing populations at these soil moistures were significantly higher at 5.0% and 4.2% day^−1^, respectively (Fig 5B). Population growth rates of growing taxa in soils at 60% and 80% field capacity were approximately twice as high than at 20% and 40% field capacity (*H* = 15.8, *p* = 0.001), with rates of 8.7% and 8.3% day^−1^, respectively (Fig 5B).

**Figure 5.**
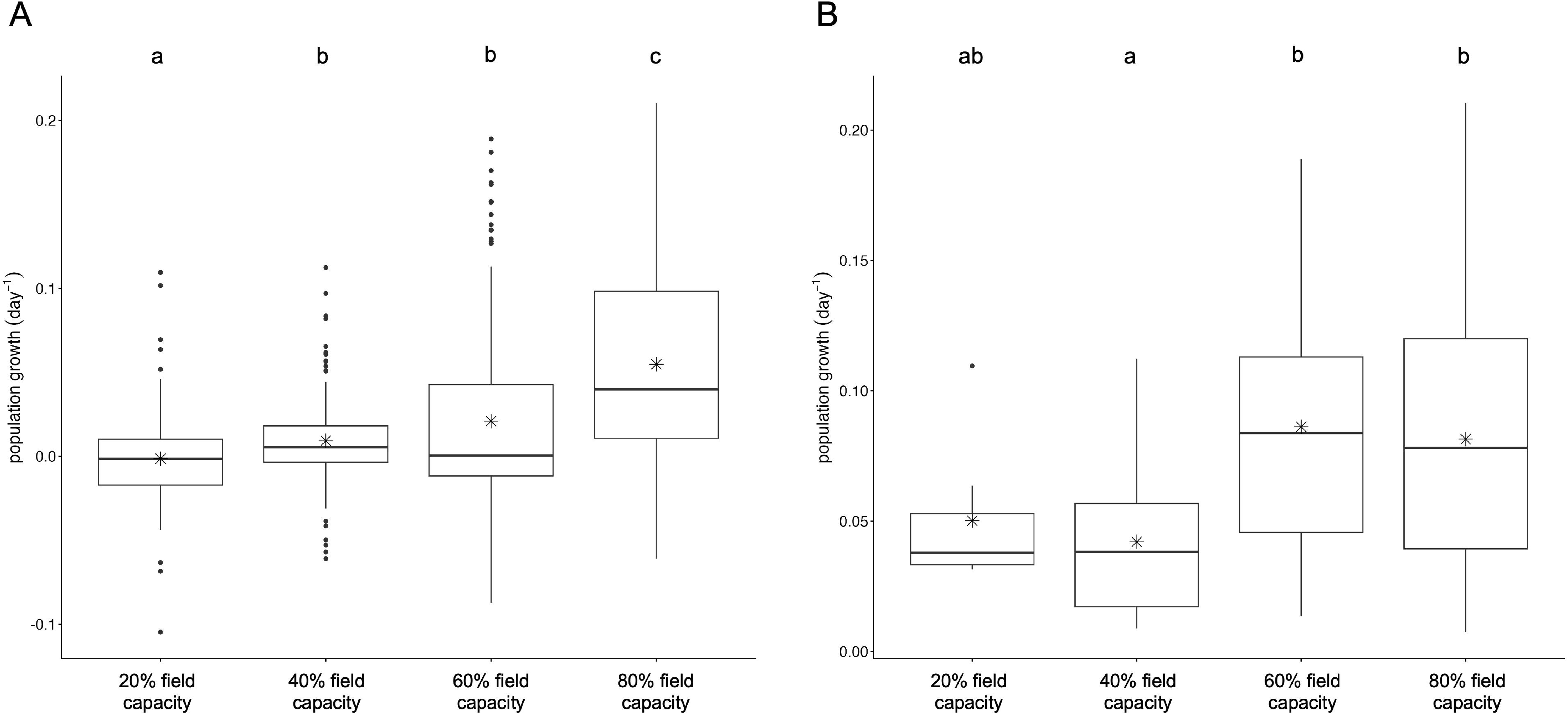
Average protist population growth rates (day^−1^) of soil incubated at four different moisture treatments: 20%, 40%, 60%, and 80% field capacity from the total protist population (A) and from the significantly growing protist population only (B). There were significant differences in population growth rates from the total protist population (*H* = 129.3, *p* < 0.0001) and from the growing protist population (*H* = 15.8, *p* = 0.001) as determined by a Kruskal-Wallis test with letters denoting significant differences from a Dunn’s post-hoc test.

Glissomonads in the Sandonidae family (TSAR;Rhizaria;Cercozoa;Filosa-Sarcomonadea;Glissomonadida) were some of the most abundant ASVs in all treatments, comprising 6.5% and 21.1% of the relative abundance of all protists in the 20% field capacity and 80% field capacity soil moisture treatments, respectively. One ASV from the Sandonidae family was growing at 20% field capacity at a rate of 3.8% day^−1^ and the richness and growth rates of ASVs from this group increased with increasing soil moisture. Four of the 26 ASVs assigned to the Sandonidae family were growing at 40% field capacity, at growth rates ranging from 0.9% to 5.7% day^−1^. In contrast, 48.9% (22 of 45) and 65.2% (30 of 46) of ASVs assigned to the Sandonidae family were growing in the 60% and 80% field capacity treatments, respectively, at average population growth rates of 9.5% day^−1^ (min 2.6% day^−1^, max 16.2% day^−1^) and 9.7% day^−1^ (min 1.8% day^−1^, max 15.9% day^−1^), respectively (Supplemental Table 1).

## Discussion

This study shows that H_2_^18^O-qSIP can measure taxon-specific DNA replication and population growth in soil protists, yielding reproducible isotope enrichment values across 18S primer sets and revealing moisture-driven changes in the growing fraction of the community. The cross-primer comparison provides methodological support for using qSIP with common 18S marker loci: enrichment values were positively related across primer datasets, and slopes moved closer to one when poorly resolved taxa were excluded. The moisture experiment demonstrates the ecological value of those quantitative measurements. Conventional 18S amplicon profiles described a community dominated by Cercozoa with relatively modest shifts in few groups, while qSIP revealed a large increase in the number, relative abundance, and growth rates of taxa making new DNA as soils became wetter.

As hypothesized, increasing soil moisture increased the richness and relative abundances of growing protist populations, where 143 populations were growing at 80% field capacity (28.8% w/w), which was dominated by ciliates and glissomonads, compared to just seven populations growing at 20% field capacity (7.2% w/w). Ciliates are some of the largest protists found in soils and as such, they require ample soil moisture for mobility and growth. For example, in an experiment where soil inoculated with *Colpoda* and *Azotobacter* was subject to suction to alter soil moisture, *Colpoda* growth was highest in saturated soils and decreased with decreasing moisture content with a cessation of growth at < 20% soil moisture despite abundant bacteria to feed on [74]. Similarly, we did not detect any ciliates growing in our lowest soil moisture treatment (7.2% w/w) while most ciliate ASVs detected in our sequencing dataset were growing in the highest soil moisture treatment (28.8% w/w).

Glissomonads are among the most abundant and diverse soil protists in molecular surveys [8, 75, 76], and our results confirm their abundance and ability to grow across conditions. Taxa in the Allapsidae family appeared more sensitive to soil moisture than Sandonidae: no Allapsidae ASVs grew at 40% field capacity, and only two of nine grew at 60%, compared to 15.4% and 48.9% of Sandonidae ASVs, respectively. This discrepancy is puzzling given their similar size (∼2.5–6.5 µM) and biflagellate morphology, though they differ in movement — Sandonidae display “jerky or jiggly” motion while Allapsidae “glide smoothly and sedately” [75] — possibly requiring different water film thicknesses. These differences may also be better resolved at finer taxonomic levels, though short amplicon sizes (<600 bp) limit species-and genus-level identification, as was the case for most ASVs here. Full-length 18S rRNA sequencing could improve resolution, though at greater costs and with ongoing reference database limitations [77].

At 20% field capacity (7.2% w/w), only 1.6% of the relative abundance of all protists were growing after a five-day incubation including two Amoebozoa, *Phalansterium* (Amoebozoa;Evosea;Variosea;Phalansteriidae) and *Flamella pleistocenica* (Amoebozoa;Evosea;Variosea;Flamellidae), as well as *Cercomonas* and *Eocercomonas* (TSAR;Rhizaria;Cercozoa;Filosa-Sarcomonadea;Cercomonadida;Cercomonadidae). *Cercomonas* are small, ameboid flagellates that glide along substrate as opposed to swimming [78], which may allow them to access prey at low soil moistures where thin water films are present, compared to swimming flagellates that would require more water filled spaces for mobility. *Flamella* are described as flattened amoebae that can form numerous fine, tapering subpseudopodia [79]. The flattened morphology of these organisms may have an advantage at low soil moistures as a survey of gymnamoebae before and after precipitation, and at depth, abundances of these shaped organisms dominated in the drier soils compared to amoebae with limax (i.e. cylindrical) morphology (5.3% – 7.3% soil moisture; [55]). While it is difficult to directly compare absolute abundances from the literature to relative abundances reported here, our estimates of the proportion of the protist population that is not growing in dry soil appear to be much higher than many reports in the literature. For example, using linear regression with data from four years of surveying gymnamoebae monthly, [54] estimated the number of encysted amoebae at soil moistures < 10% (w/w) to be 80%. Furthermore, reports of encysted amoebae ranged from 7.3% and 29.7% in surface soils and soils at 5-6cm depth, respectively, where soil moisture was 3.8% [15]. The qSIP method may underestimate the number and relative abundance of active populations by only quantifying organisms who are growing and cannot detect active organisms that are not replicating their DNA compared to counts using microscopy; however, because qSIP does not rely on rehydrating the soil in a liquid extract, we believe the results are more representative of the environmental conditions at the time of sampling.

This is the first study to our knowledge to report *in situ* population growth rates of soil protists using a molecular approach. Overall, average growth rates presented here were lower than those reported in the literature. For example, [15] counted protists during a wet-up and dry-down experiment and reported an intrinsic rate of growth of 0.15 day^−1^ during the first seven days following a wet-up, where dry soil (2.2% w/w) was wet to 20% and then dried to 7% at the time of sampling. The average growth rate of growing protist populations in our 60% field capacity soil moisture treatment, which had a similar soil moisture to the wet-up (21.4% w/w), was 0.087 day^−1^. [15] cultured their soil extracts on plates that were amended with *Klebsiella*, which may have biased the protists identified and potentially increased their growth rate estimates. They reported that their protist community was dominated by small naked amoebae (∼10 µm), making up 90-95% of the total counts, whereas we were able to detect a more taxonomically rich variety of naked amoebae, testate amoebae, ciliate, and flagellate protist populations. Using a culture independent technique may have increased our ability to detect slower growing and more cryptic taxa in comparison, thus resulting in a lower average growth rate. We also calculated population growth rates from counts reported in [19] for small flagellates (< 15 µm), large flagellates, and naked amoebae separately during a wet-up in May and dry-down in November. Population growth rates during May, when moisture increased from 2.5% to 10%, were 0.136 day^−1^, 0.125 day^−1^, and 0.193 day^−1^, respectively, for small flagellates, large flagellates, and naked amoebae, and rates increased to 0.32 day^−1^, 0.318 day^−1^, and 0.224 day^−1^, respectively, in November when soil moisture decreased from 25% to 19%. To compare, here we found average growth rates of 0.050 day^−1^ and 0.087 day^−1^ when soil moisture was 7% (similar to May sampling) and 21% (similar to November sampling), respectively. Samples from [19] were taken straight from the field during and after rain events, so soils continued to receive plant inputs, whereas we did not amend our soils prior or during our wet up, which might have resulted in slower population growth rates than in the field. This is in line with a study looking at protist population dynamics with and without wheat straw amendments, where we calculated population growth of flagellates and amoebae from reported counts and found growth to be 0.164 day^−1^ in amended soil and but no growth in the unamended soils where their numbers remained at 5 × 10^4^ g soil^−1^ [80]. Future qSIP studies that include amendments of different amounts and types could help to better elucidate bottom-up effects on protist population growth.

Together, these results show that H_2_^18^O-qSIP adds a quantitative growth dimension to protist community surveys. The method identifies growing taxa, estimates the fraction of new DNA, and translates isotope enrichment into population growth rates under an explicit growth model. In this experiment, that quantitative layer changed the ecological interpretation of the moisture gradient: less than 2% of the protist relative abundance was represented by growing taxa in dry soil (7% w/w), compared with about 60% in wet soil (29% w/w). qSIP therefore provides an important bridge between taxonomic identity and ecosystem function. The current measurements remain relative because they rely on amplicon sequencing, and qSIP specifically detects DNA replication during the incubation period. Future studies can move toward absolute population production by combining qSIP growth rates with taxon-specific 18S rRNA gene abundances from qPCR and copy-number estimates from metagenomic or single-cell approaches. That direction would allow ecologists to ask how many new gene copies or cells each taxon contributes under different environmental conditions, strengthening the link between protist identity, population growth, and soil process rates.

